# DGAT: A Dual-Graph Attention Network for Inferring Spatial Protein Landscapes from Transcriptomics

**DOI:** 10.1101/2025.07.05.662121

**Authors:** Haoyu Wang, Brittany Cody, Manuel Saavedra, Lanuza A P Faccioli, Rodrigo M Florentino, Alejandro Soto-Gutierrez, Hatice Ulku Osmanbeyoglu

**Affiliations:** Department of Biomedical Informatics, University of Pittsburgh School of Medicine, Pittsburgh, PA, USA; Department of Pathology, University of Pittsburgh School of Medicine, Pittsburgh, PA, USA

## Abstract

Spatial transcriptomics (ST) technologies provide genome-wide transcriptomic profiles in tissue context but lack direct protein-level measurements, which are critical for interpreting cellular function and microenvironmental organization. We present DGAT (Dual-Graph Attention Network), a deep learning framework that imputes spatial protein expression from transcriptomics-only ST data by learning RNA–protein relationships from spatially resolved transcriptomic and proteomic datasets. The model constructs heterogeneous graphs integrating transcriptomic, proteomic, and spatial information, encoded using graph attention networks. Task-specific decoders reconstruct mRNA and predict protein abundance from a shared latent representation. Benchmarking across public and in-house datasets demonstrates that DGAT outperforms existing methods in protein imputation accuracy. Applied to ST datasets lacking protein measurements, the framework reveals spatially distinct cell states, immune phenotypes, and tissue architectures not evident from transcriptomics alone. Here, we show that this framework accurately reconstructs spatial protein landscapes, reveals biologically meaningful tissue organization, and enables protein-level interpretation from transcriptomics-only spatial data.

## Introduction

In multicellular systems, cell function is strongly influenced by spatial context. Understanding how cells are spatially arranged, interact, and respond within their microenvironment is essential to dissect complex developmental and disease processes. Spatial transcriptomics (ST) enables mRNA profiling while preserving tissue architecture and the cellular microenvironment. Next-generation sequencing (NGS)-based ST platforms—such as the original ST^1^, 10x Visium, Slide-seq^2^, Slide-seqV2^3^, and high-definition spatial transcriptomics (HDST)^4^—use spatially barcoded probes to capture genome-wide mRNA from defined tissue regions (“spots”) that typically encompass multiple cells arranged on a two-dimensional grid.

However, transcriptomic readouts alone do not capture functional protein-level information, which is often critical for interpreting cell types/states and predicting responses to targeted therapies. Research indicates a weak correlation at the level of individual cells between the abundance of most proteins and their associated transcripts^5^. This discrepancy may arise from errors in mRNA or protein measurement, the inherent stochasticity in RNA processing, translation, and protein transport, or confounding factors such as the time delay between transcription and translation. Consequently, relying solely on mRNA expression data often fails to provide a complete picture of cellular processes and tissue architecture. Recent technological advances provide simultaneous spatial profiling of genome-wide transcript and multiplexed protein expression for up to ∼200 proteins with histopathological evaluation. These methods include Spatial PrOtein and Transcriptome Sequencing (SPOTS)^6^, Spatial Multi-Omics (SM-Omics)^7^, Spatial-CITE-seq (Cellular Indexing of Transcriptomes and Epitopes by sequencing)^8,9^, and Stereo-CITE-seq^10^ and the Visium CytAssist Spatial Gene and Protein Expression for FFPE platform by 10x Genomics. These are referred to here collectively as “spatial-CITE-seq”. Yet, due to technological barriers and cost considerations, most studies quantify the transcriptome only and do not incorporate protein measurements.

Several machine learning (ML) strategies have been developed to predict missing modalities, such as proteins, from available data. In the imaging domain, deep learning methods based on image-to-image translation have enabled virtual staining of protein markers directly from hematoxylin and eosin (H&E) images^11^. More recent advances support the prediction of multiple markers simultaneously, and tools like Multi-V-Stain improve biological relevance by optimizing all markers jointly to produce multiplexed Imaging Mass Cytometry (IMC) stains^12^.

While powerful, these image-based approaches are not applicable to transcriptomic data. In parallel, multimodal single-cell datasets have inspired methods that learn RNA– protein relationships for downstream imputation tasks^13-16^. Methods like cTP-net (single-cell Transcriptome to Protein prediction)^17^, STREAK (gene Set Testing-based Receptor abundance Estimation)^18^, sciPENN (**s**ingle **c**ell **i**mputation **P**rotein **E**mbedding **N**eural **N**etwork)^19^, scLinear^20^ and SPIDER (surface protein prediction using deep ensembles from single-cell RNA sequencing)^21^ have been tailored to learn mRNA-protein relationships from CITE-seq^5^/REAP-seq^22^ —methods that profile mRNA and surface protein expression in single cells— to generate protein estimates for scRNA-seq datasets. While these tools have shown success in single-cell settings, they are not designed to incorporate spatial structure, limiting their utility for protein inference in ST data.

Here, we present DGAT (Dual-Graph Attention Network), a computational framework for imputing spatial protein abundance from transcriptomic measurements by leveraging RNA– protein relationships learned from spatially resolved multimodal data. DGAT integrates transcriptomic profiles with spatial context using graph attention networks to infer protein expression in transcriptomics-only spatial datasets. This approach enables protein-level interpretation of spatial transcriptomics data and supports improved characterization of cellular states, tissue organization, and microenvironmental structure across diverse biological contexts.

## Results

### Overview of the DGAT framework

An overview of DGAT is shown in Fig. 1a. The framework is designed to learn from spatial CITE-seq reference datasets with paired mRNA and protein measurements. Once trained, DGAT can predict protein expression from transcript-only spatial transcriptomics (ST) datasets.

**Figure 1.**
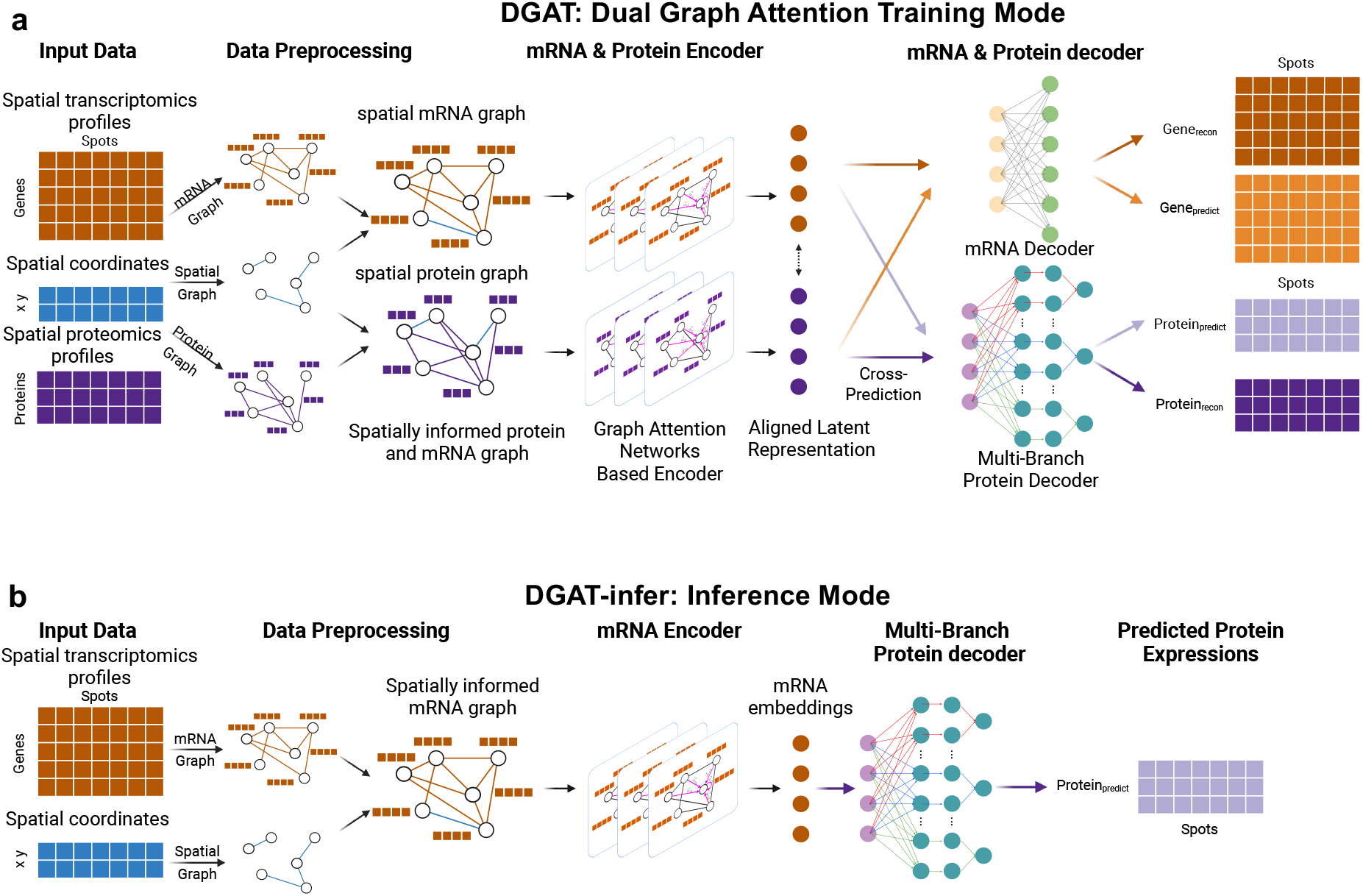
Schematic of DGAT framework. **(a)** DGAT integrates spatial transcriptomics and proteomics by constructing modality-specific heterogenous graphs based on molecular similarity and spatial proximity. Graph Attention Networks generate spatially-aware embeddings, which are aligned via latent space loss and cross-model supervision. These embeddings are decoded through a multilayer perceptron for mRNA reconstruction and a multi-branch decoder for protein prediction, enabling protein imputation from transcript-only data **(b)**.

DGAT receives as input paired gene and protein expression matrices with spatial coordinates (**Fig. 1b**). It constructs two types of graphs per sample: a spatial graph based on physical proximity, and feature similarity graphs based on transcriptomic and proteomic profiles. Each graph is processed by an independent Graph Attention Network (GAT)^23^ with feature attention layers and residual connections^24-26^ to create spatially-aware embeddings.

DGAT includes modality-specific encoders: the mRNA encoder integrates gene expression with spatial context, and the protein encoder models spatial co-expression patterns. Embeddings from both encoders are decoded in two ways: (i) a multilayer perceptron (MLP) reconstructs gene expression and (ii) a multi-branch protein decoder predicts protein levels, with each branch specialized for a distinct protein. To balance the alignment between gene and protein modalities while maintaining their distinct feature representations, we applied three types of loss functions with cross-modality supervision mechanism during training. First, a latent alignment loss encourages convergence between the transcript and protein embeddings. Second, cross-prediction losses allow each encoder to guide the opposing decoder. Finally, reconstruction losses on both modalities further constrain learning. After training, spatial protein inference can be performed by passing mRNA profiles of ST data through the pretrained mRNA encoder and protein decoder (**Fig. 1c**).

### Protein prediction benchmarking via sample holdout

We evaluated DGAT’s predictive performance using a sample-holdout strategy across six spatial CITE-seq samples, including two mesothelioma samples generated for this study and four publicly available samples (tonsil n=2, estrogen receptor-positive breast cancer, glioblastoma) (**Supplementary Table 1**). These datasets were chosen because they were among the few publicly or locally available spatial CITE-seq datasets, and they provided a diverse range of tissue types with shared protein panels, allowing systematic benchmarking of cross-sample protein prediction. Each dataset included spatially resolved transcriptomes and matched expression of ∼30 proteins (**Supplementary Table 2**) measured via the 10x Genomics CytAssist gene and protein expression platform. For each experiment, DGAT was trained on all but one sample and used to predict protein expression in the held-out sample.

Performance was assessed using the Spearman correlation coefficient (ρ) to evaluate rank concordance between predicted and observed protein levels on holdout samples, and root mean square error (RMSE) was used to quantify absolute prediction error across spatial spots. As a baseline comparison, we used normalized RNA expression values of the corresponding genes. Across all tissues, DGAT consistently outperformed the RNA baseline (**Fig. 2a, Supplementary Figure 1**). For instance, in the tonsil1 sample, the median protein-wise Spearman correlation increased from ρ = 0.308 ± 0.219 (RNA baseline) to ρ = 0.610 ± 0.230 (DGAT), while RMSE decreased from 1.61 to 0.226 (**Fig. 2a**). Similar improvements were observed across other samples (**Fig. 2a**). Notably, improvements were observed for biologically and clinically relevant markers such as CCR7, CD14, CD19, CD68, CEACAM8, CXCR5, FCGR3A, HLA-DRA, ITGAM, ITGAX, KRT5, PDCD1, PTPRC, and SDC1 (e.g. **Fig. 2b** for CD68). Consistent trends were also evident in spot-wise analyses (**Supplementary Fig. 1a**), highlighting DGAT’s ability to generalize to unseen tissue samples and improve protein inference accuracy beyond transcript-based proxies.

**Figure 2.**
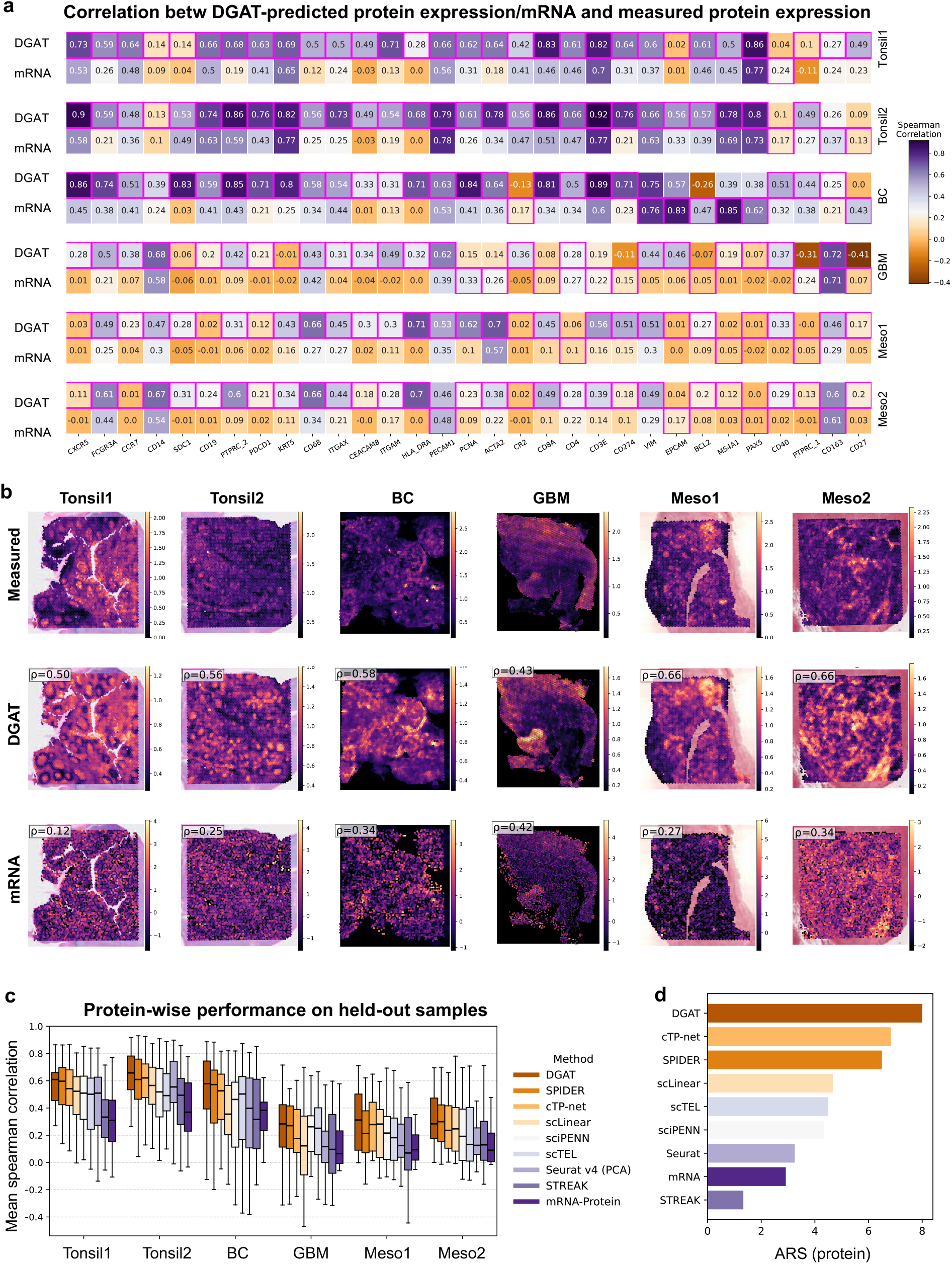
Protein prediction performance of DGAT in held-out spatial CITE-seq samples. **(a)** The heatmap shows correlations between measured protein abundances and DGAT-predicted protein abundance across different samples compared to normalized RNA expression. The highest correlation value for each protein is highlighted with a pink box. **(b)** Spatial maps of samples are shown, colored by predicted protein abundance, normalized mRNA expression, or measured protein abundance of CD68 across all samples. Correlation values between predicted protein abundance (or normalized mRNA expression) and measured protein abundance are displayed in the top-left corner of each plot. All protein and RNA expression levels (counts) are shown on a log scale. **(c)** Performance of DGAT on held-out spatial CITE-seq samples compared to mRNA, cTP-net, scLinear, sciPENN, STREAK, SPIDER, scTEL, and Seurat v4. Boxplots show Spearman correlations between predicted and actual protein expression across spots. **(d)** Barplot for ARS value of each method at the protein levels. Data are presented as the average of the six ARS values (from six datasets, six repetitions) across all experiments. Methods are ordered according to their performance.

The variation in predictive accuracy across tissues may be partly explained by the fact that our protein panel predominantly consists of immune markers, which are more abundantly and heterogeneously expressed in immune-rich tissues like tonsil and breast cancer. In contrast, glioblastoma and mesothelioma tissues typically exhibit lower immune infiltration or more complex tumor microenvironments, potentially making protein expression patterns harder to predict accurately.

Since sample-wise holdout may overestimate performance in settings where samples are highly similar (e.g., tonsil replicates; mesothelioma replicates), we additionally performed a cross-tissue (leave-one-tissue-out) evaluation. In this setting, DGAT was trained on multiple distinct tissues (e.g., breast cancer, glioblastoma, mesothelioma) and evaluated on a completely unseen tissue type (e.g., tonsil), ensuring no overlap in biological context between training and testing. As expected, absolute performance decreased relative to leave-one-sample-out evaluation. However, DGAT consistently and substantially outperformed the RNA baseline across all cross-tissue settings (**Supplementary Fig. 1b**). These results demonstrate that DGAT generalizes beyond tissues seen during training, maintaining robust accuracy despite major differences in cellular composition and tissue architecture. Together, the sample- and tissue-level evaluations confirm DGAT’s ability to infer biologically informative and tissue-relevant protein expression patterns.

We benchmarked DGAT against seven state-of-the-art models developed based on paired single cell gene expression and protein expression datasets that do not consider spatial structure: Seurat v4^27^, sciPENN^19^, cTP-net^17^, scLinear^20^, STREAK^18^, SPIDER^21^, and scTEL^28^ (**Fig. 2c, Supplementary Fig. 1a**). In all six held-out sets, DGAT achieved higher median correlation. To summarize performance across metrics, we computed an average rank score^29^ (ARS), which integrates method rankings based on correlation and RMSE. DGAT ranked highest on ARS for protein-level predictions, demonstrating robust accuracy across datasets (**Fig. 2d, Supplementary Fig. 1c**), DGAT’s advantage in spatial prediction highlights the benefits of incorporating spatial context. These results underscore DGAT’s ability to infer biologically informative and tissue-relevant protein expression patterns.

To assess whether DGAT performance varies across distinct cellular programs, we evaluated predictive accuracy stratified by canonical cell-type–associated marker proteins. Because measurements reflect mixed-cell spots, markers were grouped by dominant cell-type enrichment rather than strict cell-level assignments. Across all tissue types, DGAT consistently outperformed an mRNA-based baseline across immune, epithelial, endothelial, and stromal marker groups, with particularly strong performance for stromal and endothelial markers (**Supplementary Fig. 1d**). These results indicate that DGAT robustly captures protein expression patterns associated with diverse cellular programs, despite the mixed-cell nature of spot-based spatial transcriptomics.

### Ablation and Interpretability Analyses

To systematically assess the contributions of DGAT’s architectural components, we performed a series of ablation studies by selectively removing or modifying individual modules and design elements (**Supplementary Table 3**). First, we evaluated the impact of excluding the spatial graph, which caused a notable decline in median Spearman correlation (ρ) across datasets— from [Tonsil1: 0.610, Tonsil2: 0.658, BC: 0.579, GBM: 0.283, Meso1: 0.312, Meso2: 0.284] to [Tonsil1: 0.589, Tonsil2: 0.614, BC: 0.485, GBM: 0.153, Meso1: 0.266, Meso2: 0.178]— validating the critical role of spatial context in enhancing protein prediction. Similarly, removing the expression-based graph or relying on a single modality (e.g., RNA-only graph) further impaired performance (median ρ = [Tonsil1: 0.599, Tonsil2: 0.638, BC: 0.519, GBM: 0.228, Meso1: 0.310, Meso2: 0.291]), underscoring the necessity of multi-graph construction and modality-specific encoding.

Next, we investigated key architectural enhancements. Eliminating the feature-level attention mechanism reduced performance (median ρ = [Tonsil1: 0.585, Tonsil2: 0.661, BC: 0.533, GBM: 0.229, Meso1: 0.303, Meso2: 0.291]), highlighting its importance for prioritizing relevant molecular features. Substituting the protein-specific multi-branch decoder with a shared decoder similarly decreased accuracy (median ρ = [Tonsil1: 0.583, Tonsil2: 0.645, BC: 0.565, GBM: 0.251, Meso1: 0.305, Meso2: 0.280]), demonstrating the advantage of protein-specific mappings for flexible cross-modal learning. Additionally, ablating residual connections between GAT layers led to less stable convergence and further performance drops (median ρ = [Tonsil1: 0.571, Tonsil2: 0.622, BC: 0.460, GBM: 0.257, Meso1: 0.260, Meso2: 0.217]), suggesting these connections mitigate vanishing gradients and help preserve informative features across layers.

Finally, we examined the influence of loss functions. Removing the latent space alignment loss between RNA and protein embeddings impaired cross-modal consistency and prediction accuracy (median ρ = [Tonsil1: 0.585, Tonsil2: 0.656, BC: 0.477, GBM: 0.228, Meso1: 0.301, Meso2: 0.281]). Likewise, excluding the cross-prediction loss—where each encoder supervises the decoder of the opposite modality—resulted in reduced generalization on unseen samples

To enhance model interpretability, we performed feature importance analysis using gradient-based saliency scores^30^ on a representative tonsil sample (**Supplementary Table 4**). DGAT consistently prioritized genes with strong cell-type specificity and established protein-coding roles relevant to the predicted proteins. For instance, T cell proteins like CD3D and CD8 were associated with T cell–specific transcripts such as *GZMK, TRBC1, CXCL9*, and *SELL*, reflecting cytotoxic and memory functions. Myeloid markers CD68 and CD163 were linked to genes involved in innate immunity and myeloid differentiation, including *CXCL8, S100A8*, and *LY86*. Similarly, dendritic cell marker ITGAX (CD11c) correlated with *BATF, EBI3*, and *PLA2G7*, known mediators of dendritic cell activity. Stromal markers such as ACTA2 (α-SMA) and VIM (vimentin) were predicted using genes involved in extracellular matrix and cytoskeletal structure, including *COL14A1* and *TAGLN*. These findings demonstrate DGAT’s capacity to capture biologically meaningful RNA–protein relationships that reflect cell identity and functional states.

To further examine the biological coherence of DGAT predictions at the transcript– protein level, we analyzed the relationship between mRNA expression and DGAT-predicted protein abundance. For each protein, we computed correlations between mRNA expression and DGAT-inferred protein levels within each tissue and identified transcripts that showed consistent associations across multiple tissues (**Supplementary Fig. 2**; **Supplementary Table 5**). Across diverse protein classes, including stromal (ACTA2, VIM), endothelial (PECAM1), myeloid (CD14, ITGAX, FCGR3A), and B-cell markers (CD19, MS4A1), the most strongly associated mRNAs corresponded to well-established, cell-type–specific transcriptional programs. For example, ACTA2 and VIM were associated with extracellular matrix and contractile genes such as *COL1A1, COL1A2, DCN*, and *TAGLN*; PECAM1 correlated with endothelial-enriched transcripts including *SPARC* and *DCN*; myeloid markers were linked to canonical macrophage and dendritic cell genes such as *C1QA–C, APOE*, and *TYROBP*; and CD19-associated proteins correlated with B-cell–specific transcripts including *MS4A1, CD22*, and *CD37*. Importantly, these transcript–protein association patterns were consistently observed across tissues, indicating that DGAT captures biologically meaningful RNA–protein relationships that generalize across tissues rather than reflecting dataset-specific effects.

### DGAT enables protein inference in related spatial transcriptomics datasets

To demonstrate DGAT’s utility in ST datasets lacking protein measurements, we applied the pretrained model to publicly available human lymph node (LN) Visium data (**Fig. 3**) and triple-negative breast cancer (TNBC) tissue (**Fig. 4**) to predict the expression of 31 proteins. These tissues were selected because the DGAT training dataset includes closely related tissue types—tonsil and ER-positive (ER^+^) breast cancer—which share key structural and molecular features with LN and TNBC, respectively. This setup allowed us to evaluate the model’s capacity to generalize protein-level inference across similar yet distinct biological contexts.

**Figure 3.**
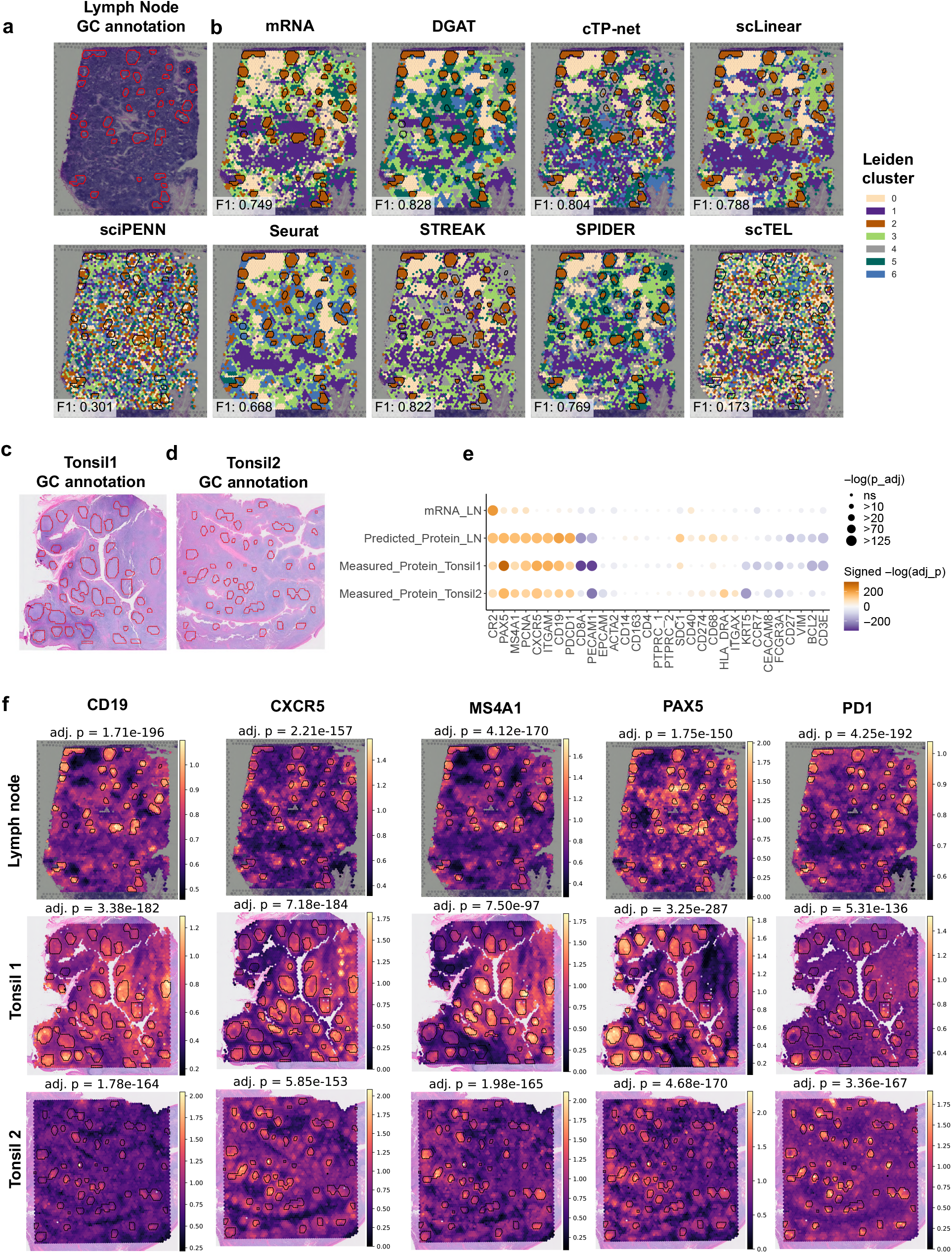
DGAT infers spatial protein expression and identifies germinal center-associated proteins in human lymph node tissue. **(a)** H&E-stained lymph node section with pathologist-annotated germinal centers (GCs). **(b)** Spatial clustering of tissue spots based on predicted protein expression from DGAT, cTP-net, scLinear, sciPENN, STREAK, SPIDER, scTEL, Seurat (anchor transfer), and raw mRNA expression, using the Leiden algorithm. DGAT achieves the highest F1 score (0.828), indicating superior spatial resolution in distinguishing GC from non-GC regions. **(c-d)** H&E-stained tonsil sections with pathologist-defined GCs used for benchmarking. **e)** Differential protein expression (GC vs. other regions) across DGAT-predicted proteins (lymph node), experimentally measured proteins (two tonsil replicates), and GC-enriched mRNA signals (mRNA_GC_LN). Circle size reflects statistical significance (−log_10_ adjusted *p*); color indicates direction of enrichment. **(f)** Spatial expression of CD19, CXCR5, MS4A1, PAX5, and PD1 in lymph node (top, DGAT-predicted) and tonsil tissues (middle and bottom, measured). DGAT recapitulates GC-localized protein expression patterns consistent with experimental data. All comparisons are statistically significant (adjusted *p* < 1×10^−15^).

**Figure 4.**
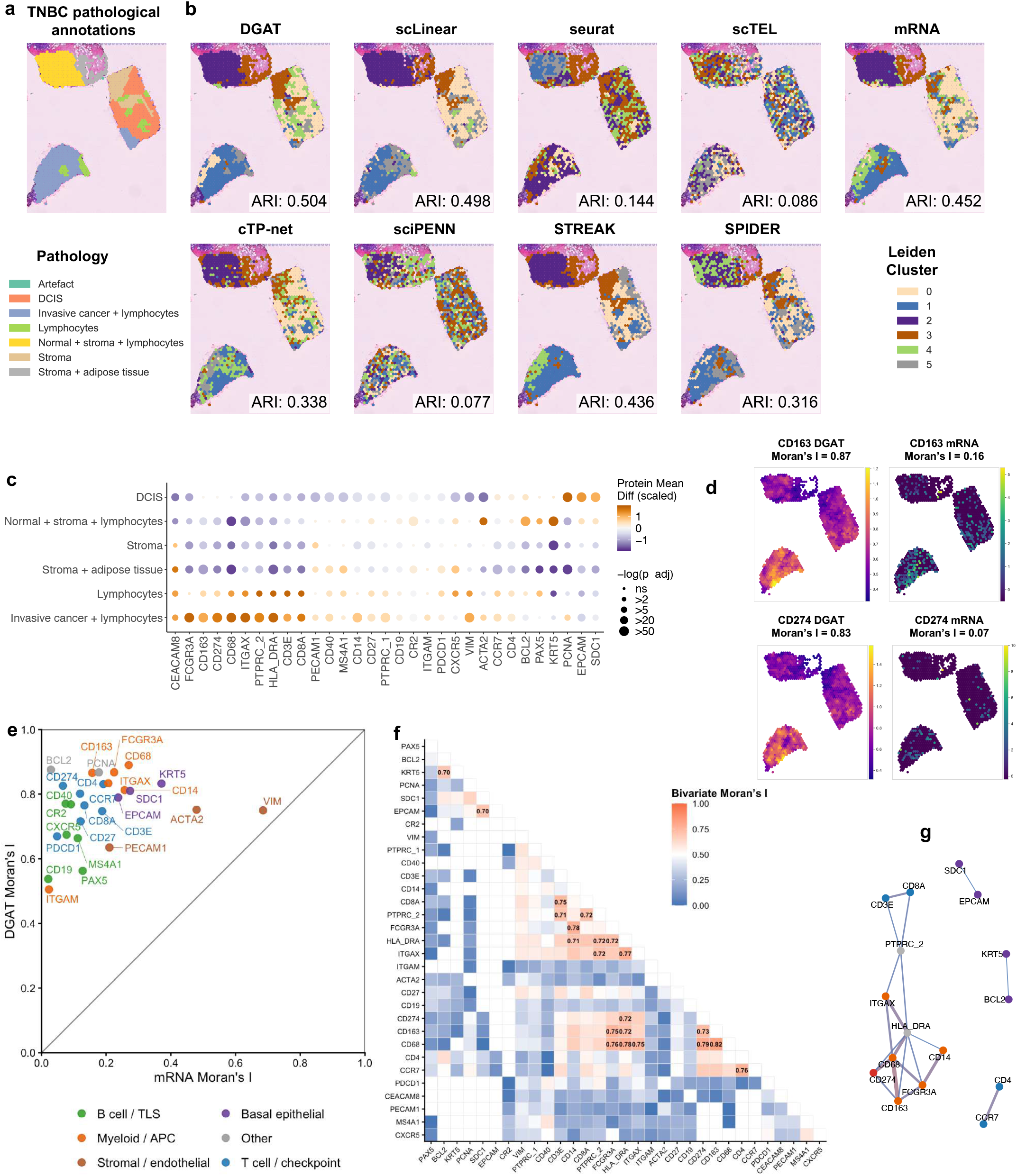
Application of DGAT to a human triple-negative breast cancer spatial transcriptomics dataset. **(a)** H&E-stained image, pathologist annotations, and **(b)** spatial clustering of a triple-negative breast cancer (TNBC) sample from Wu *et al*. based on DGAT-predicted protein expression and raw mRNA expression. Clustering performance in recovering annotated pathological regions is quantified using the adjusted Rand index (ARI). **(c)** Dot plot showing mean DGAT-predicted protein expression differences between each pathological region and all others in the sample TNBC. Dot size represents statistical significance (−log_10_ FDR) from the Wilcoxon rank-sum test. Rows and columns are hierarchically clustered. Displayed proteins are the top-ranked (lowest *P*-value) for each pathological region. **(d)** Spatial maps of representative immune markers (CD163 and CD274) showing DGAT-predicted protein expression (left) compared with corresponding mRNA expression (right). Moran’s I values quantify global spatial autocorrelation, demonstrating substantially stronger spatial coherence in DGAT-predicted protein maps relative to mRNA alone. **(e)** Comparison of global Moran’s I values for DGAT-predicted protein expression versus mRNA expression across all evaluated proteins. Each point represents a protein, colored by major functional group. Most proteins exhibit increased spatial autocorrelation following DGAT-based protein inference, indicating enhanced recovery of biologically structured spatial patterns. **(f)** Bivariate Moran’s I heatmap quantifying pairwise spatial co-localization between DGAT-predicted proteins. Warmer colors indicate stronger positive spatial associations, revealing coordinated spatial organization among immune, epithelial, and stromal protein programs. **(g)** Protein–protein spatial association network derived from significant bivariate Moran’s I interactions (I > 0.7). Nodes represent proteins, and edges denote strong spatial co-localization. Node colors indicate functional categories, including basal epithelial, myeloid/antigen-presenting cells, stromal/endothelial, and T-cell/checkpoint proteins, highlighting biologically coherent spatial neighborhoods captured by DGAT.

In the LN dataset, clustering based on DGAT-inferred protein expression revealed spatially distinct regions that closely matched histologically annotated germinal centers (GCs)^31^ (**Fig. 3a–b**). To quantitatively assess alignment with ground truth GC annotations, we framed the task as a binary classification problem, where spots assigned to clusters overlapping GCs were labeled as positive and all others as negative. This allowed us to compute classification metrics such as accuracy, precision, recall, and F1 score. DGAT-defined cluster 4 demonstrated strong alignment with GC regions, achieving an accuracy of 0.967, precision of 0.793, recall of 0.866, and an F1 score of 0.828. In contrast, transcriptome-based cluster 2 exhibited lower precision (0.613), accuracy (0.940), and F1 score (0.749), despite high recall (0.962), indicating reduced specificity and greater noise in mRNA-only clustering. These results suggest that DGAT improves both spatial resolution and biological interpretability by filtering out transcriptomic noise.

We further benchmarked DGAT against several transcriptome-to-protein prediction models that do not incorporate spatial context, including cTP-net, scLinear, sciPENN, and Seurat. Because germinal center recovery is a region-classification task, we evaluated performance using the F1 score. DGAT achieved the highest F1 score (0.828), outperforming cTP-net (0.804), scLinear (0.788), sciPENN (0.301), Seurat (0.668), scTEL (0.173), SPIDER (0.769), STREAK (0.822), and transcriptome-only clustering (0.749) (**Fig. 3b**). These results emphasize the importance of integrating spatial context to accurately reconstruct protein-level features.

To evaluate whether improved spatial domain identification could be explained solely by the cell-type specificity of the selected protein markers, we performed an additional comparison using mRNA expression restricted to the same marker gene set in these evaluation datasets. Across samples, clustering based on DGAT-predicted protein expression consistently outperformed clustering using the corresponding marker-restricted mRNA alone (**Supplementary Fig. 3**), indicating that DGAT provides added discriminatory power beyond transcriptomic representations of the same markers.

To assess the biological specificity of DGAT-inferred proteins, we evaluated their spatial enrichment in germinal center (GC) regions using the Wilcoxon rank-sum test with Benjamini– Hochberg correction. DGAT predicted protein and GC associations in the lymph node (LN) ST dataset showed strong concordance with protein–GC associations observed in two tonsil samples (Tonsil1 and Tonsil2; **Fig. 3c–d**), demonstrating the model’s ability to generalize across tissues with shared immune architecture. Canonical GC markers—including PAX5, CD19, CD20, CXCR5, CR2, CD40, PDCD1 (PD-1), and PCNA—were significantly enriched in GC regions (**Fig. 3e-f**), consistent with their roles in B cell identity, follicular homing, immune signaling, and proliferation. Notably, enrichment signals were more statistically significant using DGAT-inferred protein levels compared to mRNA expression, demonstrating DGAT’s ability to recover biologically meaningful spatial patterns obscured at the transcript level.

We next applied DGAT to spatial transcriptomics data from triple-negative breast cancer (TNBC), a subtype not explicitly included in training but biologically related through the presence of ER^+^ breast cancer samples. Clustering based on DGAT-inferred proteins again outperformed transcriptome-only clustering, achieving a higher adjusted Rand index (ARI) of 0.504 compared to 0.452, relative to expert annotations of six histopathological regions (**Fig. 4a-b**). DGAT also correctly localized canonical protein markers to their expected compartments (**Fig. 4c**). In lymphocyte-rich regions, DGAT predicted elevated expression of immune markers such as CD163, CD68, CD14, ITGAX, and FCGR3A, consistent with macrophage infiltration (**Fig. 4c**). In normal stroma and lymphocyte-enriched areas, high predicted expression of KRT5 and BCL2 reflected basal epithelial identity and anti-apoptotic signaling. In ductal carcinoma in situ (DCIS) regions, DGAT predicted increased expression of PCNA, EPCAM, and SDC1, consistent with proliferative and epithelial features characteristic of early-stage tumor progression.

To assess whether DGAT captures higher-order spatial structure beyond individual markers, we quantified spatial autocorrelation for each predicted protein. DGAT substantially increased Moran’s I relative to mRNA (**Fig. 4d–e**), revealing sharply defined spatial domains that were obscured at the transcript level. Immune markers exhibited particularly strong autocorrelation (Moran’s I = 0.75–0.86), indicating coordinated myeloid and dendritic cell localization characteristic of TNBC immune architecture.

We next interrogated protein–protein spatial organization by computing bivariate Moran’s I across all predicted protein pairs. DGAT recovered biologically coherent interaction neighborhoods, including strong co-localization between CD274 (PD-L1) and CD68 (I = 0.79), consistent with PD-L1^+^ macrophage niches; CD14 and FCGR3A (I = 0.78), reflecting monocyte–macrophage coupling; and HLA-DRA with ITGAX (I = 0.77), capturing antigen-presenting dendritic cell structures. T-cell modules were also evident, such as CCR7–CD4 (I = 0.76) marking lymphocyte homing regions and CD3E–CD8A (I = 0.75) delineating cytotoxic T-cell neighborhoods **(Fig. 4f**). These representative examples reflect well-established TNBC microenvironment programs.

A spatial protein association network constructed from high-confidence edges (bivariate Moran’s I ≥ 0.70) revealed modular organization corresponding to macrophage (CD68–CD163– FCGR3A–CD14), antigen-presenting (ITGAX–HLA-DRA), and T-cell recruitment (CCR7–CD4) communities, alongside an epithelial module (EPCAM–SDC1–KRT5) (**Fig. 4g**). Network modularity was maximized at a threshold of approximately 0.7, supporting this cutoff as balancing biological specificity and connectivity. Together, these analyses show that DGAT not only improves clustering and single-marker localization but also reconstructs functional, higher-order spatial architecture of the TNBC microenvironment, enabling richer mechanistic interpretation from transcriptomics-only ST data.

We next asked whether DGAT similarly captures spatial protein architecture in a more immune-sparse breast cancer subtype. To assess DGAT’s generalization across related breast cancer subtypes, we applied the model to an ER^+^ invasive ductal carcinoma Visium dataset. DGAT achieved the highest ARI among all baseline methods when compared with pathologist-defined tissue regions (**Supplementary Fig. 4**), indicating improved spatial domain detection from transcriptomic data alone. Beyond clustering accuracy, DGAT also recovered subtype-specific spatial protein organization, including a compact epithelial module (EPCAM–SDC1– KRT5), a modest macrophage module (CD14–CD68–CD163–FCGR3A), and a small CD3E– CD8A T-cell pair—patterns consistent with the lower immune infiltration characteristic of ER^+^ tumors (**Supplementary Fig. 4**). These findings demonstrate that DGAT preserves lineage-specific spatial structure and generalizes across biologically related breast cancer tissues, further supporting its robustness for protein inference in transcriptomics-only datasets.

### DGAT Accurately Infers Protein Expression in Unseen Tissue Types

To test DGAT’s capacity for out-of-distribution generalization, we applied the pretrained model to spatial transcriptomics datasets from human melanoma and prostate cancer—tissue types not represented in the training set. Thrane^32^ et al. manually annotated three distinct regions— melanoma, stroma, and lymphatic tissue—alongside an unannotated area (**Fig. 5a-b**). DGAT-inferred protein expression exhibited clear spatial specificity: canonical protein markers associated with each histopathological compartment were significantly enriched in their corresponding regions (**Fig. 5c**). In the melanoma region, DGAT predicted elevated expression of epithelial and proliferation markers, including KRT5, PCNA, EPCAM, and SDC1. Stromal areas showed high levels of ACTA2, PECAM1, CEACAM8, and VIM, consistent with fibrovascular and mesenchymal signatures. In lymphatic tissue, DGAT identified increased expression of immune and B/T cell markers such as CXCR5, MS4A1, PAX5, HLA-DRA, CD8, CD3, and PTPRC, reflecting the expected immune cell infiltration. At a higher level, DGAT-predicted protein co-expression patterns in melanoma revealed a modular immune architecture, with B cell and T cell markers forming tightly connected networks that are spatially segregated from epithelial and proliferative programs (**Fig. 5d**). This organization is characteristic of tertiary lymphoid structure–like immune niches observed in immunologically active melanoma.

**Figure 5.**
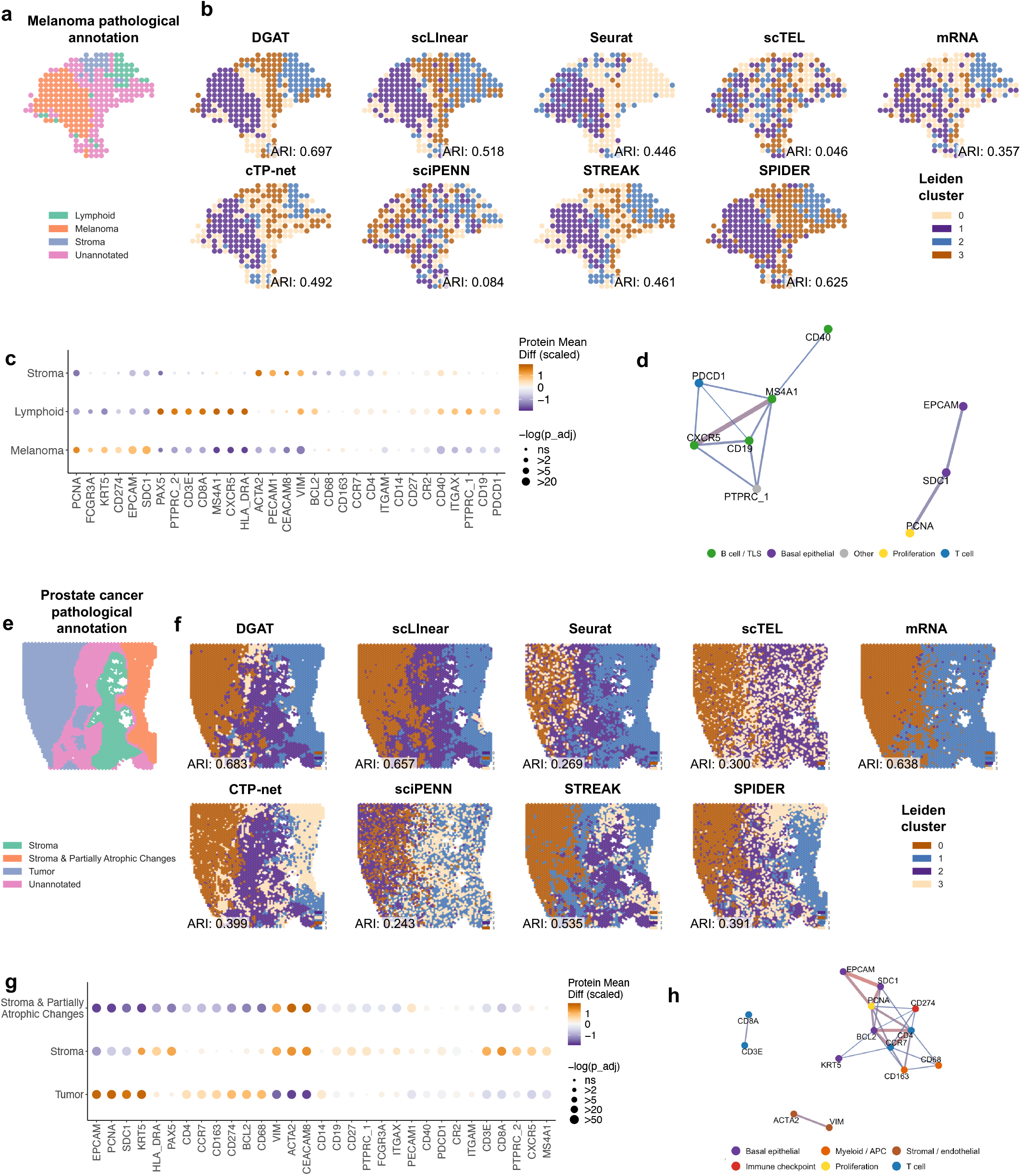
DGAT accurately infers spatial protein expression in previously unseen tissue types. **(a)** Histological annotations of the melanoma samples in the original study^32^. **(b)** Spatial clustering of melanoma tissue spots based on predicted protein expression from DGAT, cTP-net, scLinear, sciPENN, STREAK, SPIDER, scTEL, Seurat (anchor transfer), and raw mRNA expression using the Leiden algorithm. Clustering accuracy in recovering annotated pathological regions is assessed using the adjusted Rand index (ARI**). (c)** Dot plot showing mean DGAT-predicted protein expression differences between each pathological region and all others in melanoma. Dot size represents statistical significance (−log_10_ FDR) from the Wilcoxon rank-sum test; rows and columns are hierarchically clustered. **(d)** Protein–protein spatial association network in melanoma derived from significant bivariate Moran’s I interactions. Nodes represent proteins and edges denote strong spatial co-localization between DGAT-predicted protein expression patterns. Node colors indicate functional categories (B cell/TLS, basal epithelial, T cell, proliferation, and other markers), revealing biologically coherent spatial neighborhoods with a lymphoid immune module spatially segregated from an epithelial–proliferative module. **(e)** Histological annotations of the human prostate cancer samples in the original study^39^. **(f)** Spatial clustering of prostate cancer tissue spots based on protein predictions from DGAT and baseline methods, as in **b. (g)** Dot plot of DGAT-predicted protein expression differences between pathological regions in prostate cancer, with statistical significance and clustering visualized as in **f. (h)** Protein–protein spatial association network in prostate cancer derived from significant bivariate Moran’s *I* interactions (*I* > 0.7). Nodes represent proteins, and edges indicate strong spatial co-localization, with edge thickness proportional to association strength. Node colors denote functional categories, including basal epithelial, myeloid/antigen-presenting, stromal/endothelial, proliferation, immune checkpoint, and T-cell markers. The network exhibits a modular organization comprising an epithelial–proliferative module (EPCAM–SDC1–PCNA– KRT5), a myeloid/immune checkpoint–associated module (CD68–CD163–CD274), a T-cell module (CD3E–CD8A), and a stromal module (ACTA2–VIM), highlighting biologically coherent spatial architecture captured by DGAT.

DGAT also generalized effectively to prostate cancer spatial transcriptomics data, recovering spatially distinct protein expression patterns corresponding to epithelial, stromal, and immune-rich compartments (**Fig. 5e–f**). In this dataset, DGAT achieved the highest adjusted Rand index (ARI) of 0.683, outperforming all baseline methods in accurately recapitulating pathologist-defined tumor, stromal, and partially atrophic regions. The mRNA-only model performed moderately (ARI 0.638), while scTEL, cTP-net, STREAK, SPIDER, Seurat, and sciPENN underperformed with ARIs of 0.3, 0.399, 0.535, 0.391, 0.269, and 0.243, respectively. Within prostate cancer tissue, tumor regions were specifically enriched for CD63 and CD9, along with epithelial markers EPCAM and VIM (**Fig. 5g**). Stromal compartments were enriched for immune-related proteins CD14 and CD68, consistent with macrophage and monocyte infiltration. Partially atrophic areas exhibited intermediate expression profiles, reflecting a mixture of epithelial and immune signals indicative of a transitional microenvironment. In contrast to melanoma, DGAT-predicted protein networks in prostate cancer were denser and more interconnected, with strong coupling between epithelial, myeloid, and immune checkpoint– associated markers (**Fig. 5h**). This pattern is consistent with a more immunosuppressive and heterogeneous tumor microenvironment, characteristic of prostate cancer.

Across both datasets, DGAT-predicted protein landscapes closely mirrored known histological boundaries despite the domain shift from training tissues. These results suggest that DGAT captures broadly conserved RNA–protein relationships that remain informative across diverse tissue contexts. Importantly, DGAT’s ability to recover biologically meaningful protein profiles in the absence of paired proteomic data underscores its potential as a powerful tool for retrospective spatial transcriptomics analysis, cross-tissue protein mapping, and hypothesis generation in under-characterized disease settings. Together, these results demonstrate that DGAT not only generalizes spatially across unseen tissue types, but also preserves disease-specific protein organization and microenvironmental structure

### Experimental validation of DGAT across independent tissue types and disease contexts

To evaluate the generalizability of DGAT beyond the tissue types used for training, we applied the pretrained model to five independent in-house spatial transcriptomic datasets encompassing three kidney samples (two with chronic kidney disease [CKD] and one normal kidney) and two lung samples from chronic lung disease (CLD) (**Supplementary Table 1**). These datasets represent distinct organ systems and disease contexts not included during model training, providing a robust assessment of DGAT’s transferability to unseen tissues.

Predicted protein-presence distributions derived from the ST sections were compared with immunofluorescence (IF) staining of adjacent tissue sections for three representative proteins—ACTA2 (αSMA), PECAM1/CD31, and VIM—selected to represent smooth muscle, endothelial, and mesenchymal compartments, respectively. DGAT-imputed protein-presence (**Fig. 6a**) closely recapitulated spatial IF patterns across all samples, accurately capturing key tissue structures such as vascular networks (PECAM1/CD31), smooth muscle regions (ACTA2/αSMA), and stromal domains (VIM).

**Figure 6.**
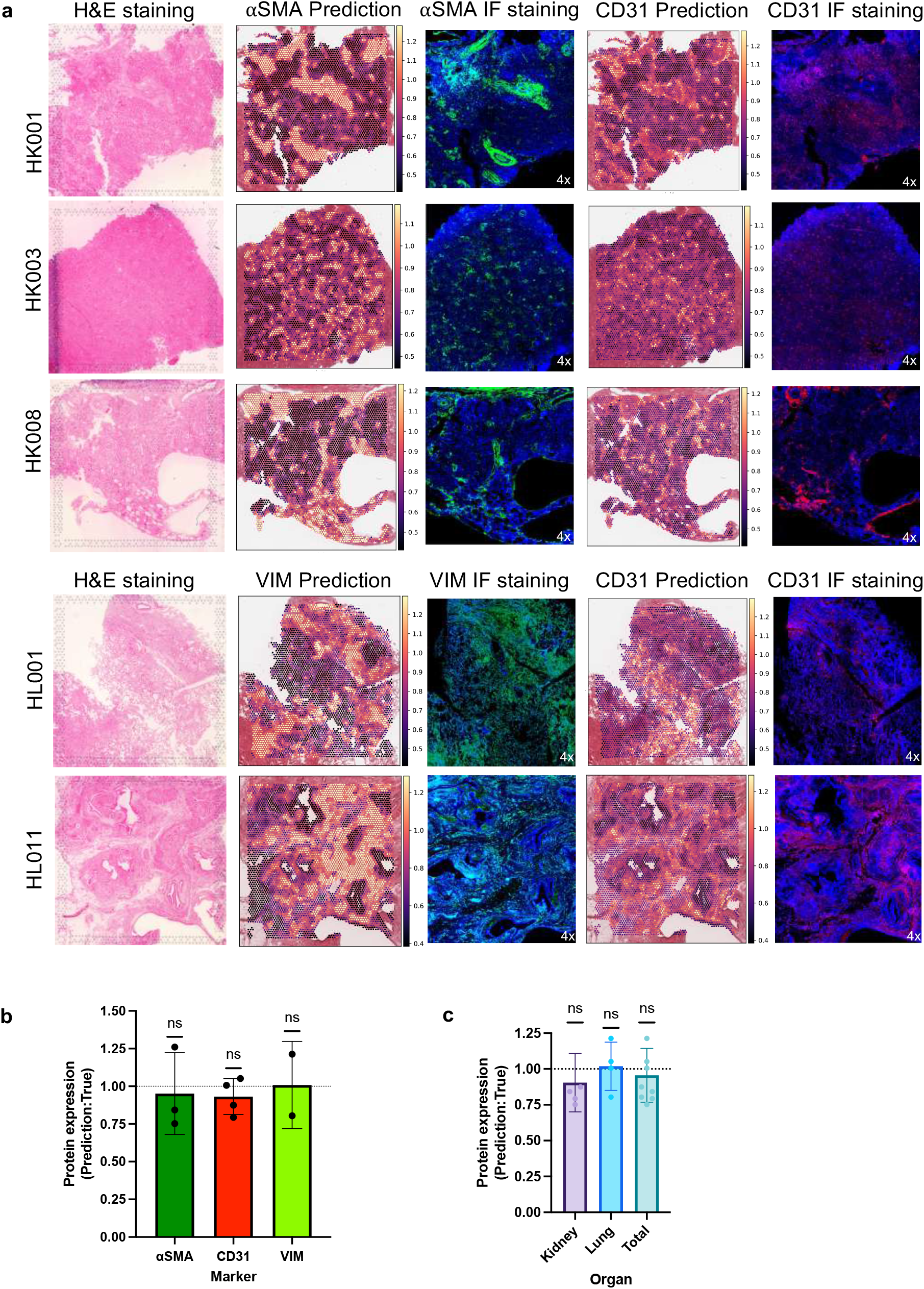
Immunofluorescence validation of DGAT-imputed protein expression across independent tissue types and disease contexts. **(a)** Representative images comparing hematoxylin and eosin (H&E) staining (from the same tissue block used for spatial transcriptomic profiling), DGAT-imputed protein spatial maps, and corresponding immunofluorescence (IF) staining. IF was performed on adjacent tissue sections from the same anatomical regions. αSMA (ACTA2) and VIM are shown in green, CD31 (PECAM1) in red, and nuclei are counterstained with DAPI (blue). Kidney samples include HK001 (normal), HK003 and HK008 (chronic kidney disease, CKD); lung samples include HL001 and HL011 (chronic lung disease, CLD). **(b)** Ratios of predicted-to-measured expression area, averaged across three manually selected regions of interest (ROIs) per sample and stratified by marker (αSMA, CD31, VIM). **(c)** Predicted-to-measured expression area ratios grouped by organ (kidney and lung) and across all samples. The dashed line indicates the expected ratio of 1. One-sample *t*-tests were performed against the null hypothesis (ratio = 1), revealing no systematic bias (ns). *P* values are reported in the panel. Error bars represent the standard error of the mean (SEM).

Quantitative analysis further confirmed the fidelity of DGAT predictions. The ratios between predicted and IF-detected protein-positive areas were not significantly different from unity (one-sample t-test, p > 0.05 across all markers and organs; **Fig. 6b-c**), indicating no systematic over-or underestimation of protein presence. Together, these findings demonstrate that DGAT achieves robust, biologically consistent protein predictions across independent organ systems and disease contexts, underscoring its generalizability beyond the datasets used for training.

## Discussion

The ability to infer spatial protein expression from transcriptomic data alone opens new avenues for understanding tissue organization, cellular heterogeneity, and microenvironmental dynamics in health and disease. Here, we introduced DGAT, a deep learning framework trained on spatial CITE-seq data that accurately imputes spatial protein expression in transcriptomics-only datasets. Our results demonstrate that DGAT not only recovers biologically meaningful protein distributions but also enhances key downstream analyses such as spatial clustering and spatial domain identification (e.g. GCs). Further, DGAT’s application to in-house malignant pleural mesothelioma tissue, a rare and understudied cancer^33,34^, demonstrates its value in expanding spatial proteomic profiling beyond well-characterized diseases. Hence, DGAT enables proteome-level insights in datasets where protein measurements are unavailable, offering a computational bridge between transcriptomic accessibility and proteomic relevance.

A major advantage of DGAT lies in its ability to generalize across tissue types. We showed that a model trained on diverse spatial CITE-seq datasets could be effectively applied to unrelated transcriptomics-only datasets, including lymph node and breast cancer tissues, where it improved the concordance between predicted spatial clusters and expert pathological annotations. This suggests that the protein–transcript relationships learned by DGAT are sufficiently robust to capture conserved biological patterns, even across different microenvironments.

Building on these results, we further validated DGAT using additional in-house ST datasets from chronic kidney disease, a normal kidney, and chronic lung disease (CLD). DGAT-predicted protein distributions showed strong spatial concordance with experimentally measured immunofluorescence signals for ACTA2 (αSMA), PECAM1/CD31, and VIM, confirming its robustness across independent tissue types and pathophysiological contexts.

While DGAT demonstrates strong generalization across tissues profiled here, several limitations warrant consideration. First, DGAT’s protein decoder is configured for a fixed protein panel and therefore requires retraining when applied to datasets with different protein targets, although retraining remains computationally efficient. Second, DGAT operates at the resolution of spatial transcriptomic spots, which may contain mixtures of cell types. Finally, DGAT relies on the availability of paired spatial transcriptomic–proteomic reference datasets for supervised training, which are currently limited in number. As additional multimodal spatial datasets become available, we anticipate that DGAT’s performance and applicability will further improve. Moreover, because mRNA–protein relationships can vary substantially by tissue type and cell state, models trained on immune-rich or cancer samples may require retraining or fine-tuning to achieve optimal performance in tissues with distinct molecular landscapes, such as hepatic, neural, or fibrotic tissues. Similarly, while we evaluated DGAT on approximately 30 immune-related proteins, scaling to larger panels measured in some CITE-seq studies (200–300 proteins) should be feasible provided these markers are well represented in the training data, though this remains to be systematically benchmarked.

Regarding training data availability, DGAT was trained and evaluated on all high-quality Visium CytAssist Spatial Gene + Protein Expression datasets that were publicly accessible or generated in-house at the time of analysis, including tonsil, breast cancer, glioblastoma, and malignant mesothelioma. The framework is tailored to spot-based spatial transcriptomic– proteomic assays that provide co-registered gene and protein measurements. Although imaging-based proteomic platforms such as CODEX, CyTOF, and multiplexed immunofluorescence offer extensive protein coverage, they currently lack matched spatial transcriptomes required for supervised training. As multimodal datasets integrating spatially resolved transcriptomes and proteomes across additional tissues become available, DGAT can be retrained or fine-tuned to expand its applicability. Broadening the diversity of training data will be essential for capturing tissue-specific mRNA–protein relationships and further strengthening the model’s generalizability.

Current spatial transcriptomics and spatial CITE-seq platforms are limited by their resolution and coverage. Capture spots often encompass multiple cells, leading to mixed signals that obscure cell-specific expression profiles. These technical constraints pose challenges for fine-grained spatial inference, particularly in complex tissues like tumors. However, emerging technologies—such as Visium HD and other high-resolution spatial omics platforms—promise to address these limitations by offering near-single-cell resolution and expanded molecular profiling. We anticipate that DGAT will be well suited for these next-generation datasets, where finer spatial granularity may further improve the fidelity of protein inference.

Looking forward, several extensions of this work are possible. First, expanding the training corpus with additional spatial CITE-seq datasets from diverse tissues and disease contexts could improve the model’s generalization capacity and resilience to dataset-specific biases. Second, integrating histopathological images (e.g., H&E or multiplexed immunofluorescence) may allow DGAT to incorporate morphological context, further enhancing protein prediction accuracy and interpretability. Lastly, beyond clustering, DGAT could support a broader range of spatial analyses, including ligand–receptor interaction mapping, cell–cell communication inference, and spatial trajectory reconstruction, offering a powerful toolset for dissecting tissue biology and informing therapeutic discovery.

In conclusion, DGAT provides a scalable, data-efficient solution for inferring protein-level spatial maps from transcriptomic inputs alone. As spatial omics technologies continue to evolve, DGAT stands to facilitate deeper biological insights in both basic and translational research settings—particularly where proteomic measurements are unavailable or impractical. DGAT represents a step toward democratizing spatial proteomics, enabling researchers to unlock functional insights from transcriptomic-only data without requiring expensive or technically demanding protein assays.

## Methods

### Malignant mesothelioma Visium CytAssist v2 library preparation and sequencing

Two formalin-fixed & paraffin-embedded (FFPE) malignant mesothelioma tissue blocks were obtained from National Meosthelioma Virtual Bank. Malignant mesothelioma samples were assessed for RNA quality, and all samples had a DV200 score above the minimum threshold ≥30%. Spatial transcriptomic and proteomic libraries were generated with Visium CytAssist for FFPE v2 gene expression kit (10x Genomics: 1000520) with the Human Immune Cell Profiling Panel (10x Genomics: PN-1000607). 5 micron FFPE sections were placed on Schott Nexterion Hydrogel Coated Slides (Schott North America: 1800434) or Fisherbrand Superfrost Plus Slides (Fisher: 22-037-246) and processed through the 10x CytAssist for FFPE v2 protocol according to the manufacturer’s instructions. H&E images were taken at 10x magnification using an EVOS 7000 Imaging System (Thermo Fisher Scientific).

Library QC was completed with an Agilent TapeStation 4150. Libraries were normalized and pooled to 2nM prior to loading on an Illumina NextSeq 2000, using a P3 100 flow cell with a target of 125 million reads per sample for transcriptomic libraries and 25 million reads per sample for proteomic libraries.

### Kidney and lung Visium v2 library preparation and sequencing

To assess DGAT generalizability, six in-house FFPE samples were profiled using the Visium v2 Spatial Gene Expression platform (10x Genomics). This cohort included four kidney tissues (two with chronic kidney disease and one normal) and two lung tissues from chronic lung disease (**Supplementary Table 1**). Samples were assessed for RNA quality, and all FFPE samples had a DV200 score above the minimum threshold of ≥30%. Spatial transcriptomic libraries were generated with the Visium v2 Human Transcriptome gene expression kit (10x Genomics: 1000443). Five-micron FFPE sections were placed on Superfrost Plus slides (FisherScientific) and processed through the 10x Visium v2 for FFPE protocol according to the manufacturer’s instructions (CG000518, CG000520, CG000495). H&E images were taken at 20x magnification using a Leica Aperio CS2 slide scanner (Leica Biosystems). Library QC was completed with an Agilent TapeStation 4150. Libraries were normalized and pooled to 2nM prior to loading on an Illumina NextSeq 2000, using a P4 100 flow cell with a target of a minimum of 120 million reads per sample for transcriptomic libraries.

### Data Preprocessing

We used datasets from three different platforms, including Visium CytAssist Gene and Protein Expression (tonsil1, tonsil2, breast cancer, glioblastoma from 10x Genomics and in-house malignant mesothelioma1, malignant mesothelioma2), 10x Visium platform (human breast cancer and human prostate cancer), the ST platform (human melanoma^32^). A detailed description and data sources are provided in **Supplementary Table 1**.

For all in house and public spatial CITE-seq (Visium CytAssist) and spatial transcriptomics datasets, low-quality spatial barcodes (spots) were removed if they contained fewer than 700 gene expressed or had >30% of UMIs derived from mitochondrial transcripts. Genes not expressed in at least 2.5% of spots were excluded to reduce technical noise and sparsity. Gene expression matrices were normalized to 10,000 UMIs per spot, log-transformed, and scaled to unit variance per gene. Values were clipped to a maximum of 10 to mitigate the influence of outliers.

Protein expression (ADT counts) was normalized using the centered log-ratio (CLR) method. After preprocessing, the dataset comprised 11,535 genes and 31 proteins across six samples, which were integrated via shared features into a unified dataset for downstream modelling. The SCANPY toolkit^35^ was utilized for all preprocessing steps.

### Model Architecture: DGAT (Dual Graph Attention Network)

#### Graph Construction

We denoted the mRNA and protein expression matrices as X_mRNA_ ∈ ℝ^N×G^ and X_protein_ ∈ ℝ^N×P^, where N was the number of spots, and G and P were the number of genes and proteins, respectively. Each spot was modeled as a node in a graph with features defined by either mRNA or protein expression. Two types of adjacency matrices were constructed: spatial adjacency (**A**_spatial_) and modality-specific adjacency (**A**_mRNA_, **A**_protein_). The spatial graph connects each spot to its six nearest physical neighbors based on Euclidean distance of spot coordinates. Molecular graphs were built using k-nearest-neighbor (k=10) graphs in principal component (PCA) space (retaining 85% of variance) for each modality. The final adjacency matrices used for each modality were defined as:

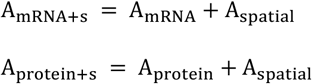

#### Encoder design

While the mRNA and protein encoders differ in their input dimensionalities— reflecting the distinct numbers of mRNA and protein features—their internal architectures are otherwise identical to ensure consistent feature extraction across modalities. Each encoder consists of three Graph Attention (GAT) layers, interleaved with Layer Normalization and skip connections, enabling the learning of robust, generalized latent representations. The inputs to each encoder are (X_mRNA_, A_mRNA+s_) or (X_protein_, A_protein+s_). The encoding process is defined as:

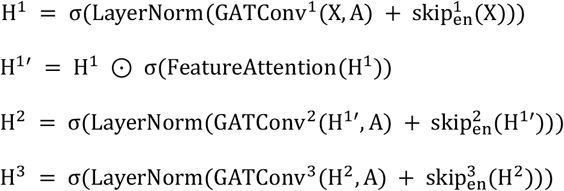

where σ is the LeakyReLU activation, and ⊙ denotes element-wise multiplication. Multi-head feature attention is applied as:

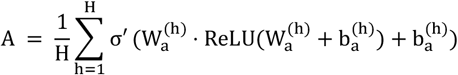

where H denotes the total number of attention heads, W_a_^(h)^ and b_a_^(h)^ are the learnable weight matrices and bias terms for attention head *h*, respectively, and σ’ is the sigmoid activation function used for gating.

The output of each encoder is a modality-specific latent representation computed as:

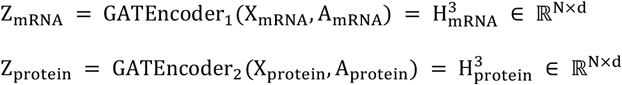

where d=1024 is the latent dimensionality and N is the number of spots or nodes in the graph. These latent embeddings are subsequently passed to the decoder modules for downstream tasks.

#### Decoder design

In the DGAT model, decoder modules are tasked with reconstructing mRNA and protein expression from the learned latent embeddings with distinct architecture design. The mRNA decoder is implemented as a fully connected feedforward neural network with residual connections, designed to map latent embeddings back to the original mRNA feature space:

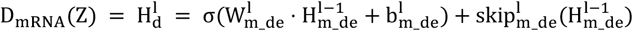

where 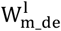 denotes learnable weights for l-th layer in mRNA decoder. 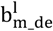 is learnable bias term. σ represents LeakyReLU activation. 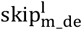 is residual (skip connection) projection layer. 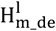 is the layer output of the mRNA decoder.

The protein decoder consists of a shared two-layer trunk followed by multiple independent branches, one per protein, to model protein-specific reconstructions:

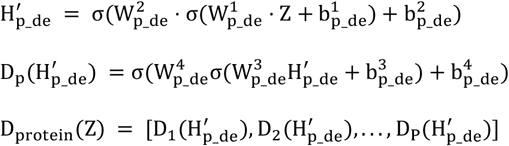

where 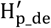 denotes the shared feature extracted by the trunk in the protein decoder. 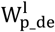 represents the learnable weights for l-th layer. 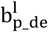 is learnable bias term. D_p_ is a protein branch which predicts one protein’s expression. D_protein_ is formed by concatenating the outputs of each protein-specific branch.

#### Loss functions

We designed three types of loss functions, each serving a distinct purpose to facilitate mRNA-to-protein prediction. The alignment loss defined as ℒ_align_ = MSE(Z_mRNA_, Z_protein_), where MSE is mean square error, aims to maximize the concordance between the mRNA and protein latent embedding spaces. The reconstruction loss, defined as

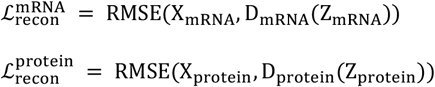

Where RMSE represents the root mean square error, ensures that the latent representations produced by each encoder retain its distinct modality-specific features. Additionally, we introduce a prediction loss by cross-feeding the latent embeddings into the decoder of the opposite modality, defined as

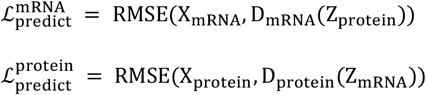

supervises and reinforces the model’s ability to perform cross-modality inference. Finally, the total loss function is defined as:

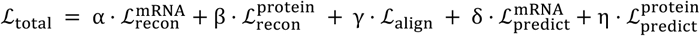

where α, β, γ, δ, η are hyperparameters which control the relative importance of each loss component.

#### Model architecture and hyperparameters

DGAT comprises two three-layer Graph Attention Network encoders (GATEncoder) followed by two parallel decoders for mRNA and protein prediction. The GATEncoder first applies a GATConv layer with an output dimension of 2048, combined with a residual linear projection from input dimension to 2048, with a layernorm, LeakyReLU (negative slope = 0.2) and dropout (p = 0.4) after each graph attention block. After the graph attention block, DGAT utilizes a multi-head channel-wise feature attention mechanism with 16 independent attention heads to automatically assign to improve the aggregation of contextual information. Each head is a two-layer gating transformation (2048 to 512, ReLU, Linear 512 to 2048, Sigmoid), and the averaged gates are used to reweight features. The second encoder block consists of a GATConv layer with hidden size 1024, paired with a residual linear projection (2048 to 1024), LayerNorm, LeakyReLU(0.2), and dropout (p = 0.4).

The third block reduces the representation to the latent dimension (1024) using a GATConv (1024 to the hidden layer dimension) and a residual projection, again followed by LayerNorm, LeakyReLU(0.2), and dropout (p = 0.4). The protein decoder consists of a shared trunk and protein-specific branches. The trunk maps hidden layers to 512 and then 256 with LayerNorm and LeakyReLU (0.2). Each protein branch is a small multi-layer perceptron (256 to 64, LayerNorm, LeakyReLU(0.2), 64 to 1), inferring a protein expression. The mRNA decoder is a three-layer feed-forward network with residual connections, where hidden layer was projected to 512, then to 1024, finally back to the mRNA dimension.

#### Model training

The training and evaluation process for the DGAT model is designed to ensure the accurate cross-modality prediction for protein expression from spatial transcriptomics data. The key steps are as follows: The weights α, β, γ, δ, η were set based on preliminary experiments to *α* = 5, *β* = 1, *γ* = 1, *δ* = 3, *η* = 5. The alignment loss directly couples the mRNA and protein latent spaces, which can lead to over-regularization if overemphasized. To enhance the model’s generalization, we applied dropout rate of 0.3 for both mRNA encoder and protein encoder. Moreover, to stabilize the training and avoid over-optimization for the alignment loss, the weight of it is set to 0 if the loss is lower than 0.015. Limiting it dynamically retains modality-specific nuances, reducing the risk of both embeddings collapsing into a single, less informative and less biological meaningful space. Separate Adam optimizers are used for each encoder and decoder. The learning rates for encoders are 5 × 10^−4^, while they are 1 × 10^−4^ for decoders. In addition, gradient norms are clipped to 1.0 to prevent exploding. In addition, A small weight decay of 2 × 10^−5^ is applied to the encoders to improve generalization. The learning rate is reduced by 20% every 10 epochs to encourage convergence. Finally, training is stopped if the total loss does not improve for 10 consecutive epochs.

We adopt a noise-based early stopping criterion (“EB-criterion”) that does not require held-out validation data^36^. The key idea is to monitor when the model’s parameter updates become dominated by noise rather than useful learning signals. To do this, we estimate the signal-to-noise ratio of parameter gradients during training. When this ratio becomes low— indicating that continued training is unlikely to yield meaningful improvement—we stop training. To ensure robustness, we delay the stopping check until after 50 epochs and smooth the evidence signal by averaging over the most recent 10 epochs. Training stops when this averaged signal crosses a predefined threshold (0.96), indicating that learning has effectively plateaued.

#### Model performance

Leave-One-Out Testing (LOOT) is used to evaluate the model’s generalization. Each sample in the dataset is treated as a test set while the remaining samples are used for training. This ensures robust performance evaluation across diverse tissue types. To evaluate the accuracy of the protein imputation methods, we calculated the following metrics: (a) Spearman correlation coefficient (ρ): This quantified the rank correlation between the predicted and measured protein expression levels, helping us assess how well the imputed data preserves the ordering of the original data. (b) Root Mean Square Error (RMSE): This metric measured the overall error between imputed and actual protein levels, providing a measure of model accuracy. (c) Average Rank Score (ARS): We computed ARS as a combined metric (ref), incorporating both Spearman correlation and RMSE. This gave us an overall performance indicator, with a higher ARS value representing better accuracy across the evaluation metrics. These metrics were used to assess the accuracy of protein imputation at both the protein level (quantifying the accuracy of protein expression values) and the spot level (assessing accuracy for individual spots).

#### Computational efficiency

DGAT’s decoder uses a fixed set of protein-specific output branches; therefore, retraining is required when users modify the protein panel. To quantify computational requirements, we measured DGAT’s training time across datasets containing one to six spatial CITE-seq samples on a single NVIDIA L40S GPU (**Supplementary Table 6**). Training times scaled moderately with dataset size, ranging from 49.1 s (1 sample) to 401.4 s (6 samples), with an average of ∼55–66 epochs. Owing to its lightweight architecture, DGAT remains computationally efficient to retrain, and full model retraining can be completed within minutes even on modest GPU hardware. Future work will explore decoder designs that flexibly accommodate variable protein panels without retraining.

### Benchmarking procedure

We compared DGAT against cTP-net^17^, sciPENN^19^, scLinear^20^, Seurat, STREAK^18^, SPIDER^21^, and scTEL^28^. All baseline models were retrained on the same spatial CITE-seq datasets used for DGAT, and pretrained scRNA-seq–derived weights were not used. Each method was implemented following the codebase and documentation provided in the respective publications.

#### Model-specific implementations

For cTP-net, protein abundances were inferred from gene expression matrices using the official training script with default parameters. SAVER-X denoising was omitted due to runtime and software dependency constraints. For sciPENN, preprocessing and model training were conducted using the default parameters provided in the GitHub repository (https://github.com/jlakkis/sciPENN). For Seurat (v4), data were processed using SCTransform, dimensionality reduction was performed with PCA, and protein imputation was carried out using FindTransferAnchors and TransferData with default settings (https://satijalab.org/seurat/). For scLinear, we followed the tutorial provided on Zenodo (https://zenodo.org/records/10602824) using default settings. For STREAK, we followed the tutorial provided in the GitHub repository (https://github.com/azkajavaid/STREAK). For scTEL, we followed the tutorial provided in the GitHub repository (https://github.com/142857cyy/scTEL). For SPIDER, we followed the tutorial provided in the GitHub repository (https://github.com/Bin-Chen-Lab/spider/tree/main).

### Cell-type–stratified performance analysis

To assess model performance across distinct cellular programs, marker proteins were grouped by their canonical cell-type enrichment (T cell, B cell, myeloid, antigen-presenting, epithelial, endothelial, stromal, proliferative, immune checkpoint, and leukocyte). For each marker, DGAT-predicted protein levels were correlated with measured protein expression across all spatial spots using Spearman correlation, and correlation coefficients were summarized within each marker group. Because measurements reflect mixed-cell spots, markers were interpreted as cell-type-enriched rather than cell-type-specific. An mRNA-based correlation model served as the baseline for assessing DGAT’s predictive improvements.

### Feature importance analysis

To interpret the model’s protein prediction behavior and identify key transcriptomic contributors, we performed a feature importance analysis based on gradient attribution. For each protein-specific decoder branch, we computed the gene-level importance scores by aggregating the input-gradient product 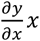 across all cells, normalized by z-scoring. This method highlights genes whose expression changes have the strongest influence on protein prediction, thereby capturing both sensitivity and directionality.

### Spatial autocorrelation and protein association analysis

To assess the spatial organization of predicted protein expression, we quantified spatial autocorrelation using Moran’s I^37^, a standard measure of spatial dependence. For each protein, Moran’s I was computed across spatial spots using DGAT-predicted protein levels and a spatial weight matrix defined based on spot adjacency. Moran’s I values were compared with those computed using mRNA expression of the corresponding genes to assess gains in spatial coherence attributable to DGAT.

To evaluate higher-order spatial relationships between proteins, bivariate Moran’s I was computed for all pairs of predicted proteins, measuring spatial co-enrichment between protein *i* at a given spot and protein *j* in neighboring spots. Protein association networks were constructed by thresholding bivariate Moran’s I values to retain high-confidence spatial associations. Nodes represent proteins and edges represent significant spatial co-localization. Network modularity was assessed across a range of thresholds to identify a cutoff that balanced biological specificity and network connectivity.

### Immunostaining and quantification

Human kidney, and lung tissue were fixed in 4% paraformaldehyde for 12 hours and 70% ethanol overnight at 4°C and then embedded in paraffin. Five-micron sections were mounted on glass slides for immunofluorescence (IF). Adjacent sections were used to assess protein presence by IF to validate DGAT predictions made on the spatial transcriptomics slides. Each sample was first stained with hematoxylin and eosin for histological examination. Slides were deparaffinized with xylenes and dehydrated with ethanol. Antigen unmasking was performed by boiling in 10 mM citrate buffer, pH 6.0. The slides were then blocked for 1 hour with 10% donkey serum, left incubating overnight at 4oC with primary antibodies listed in the **Supplementary Table 7**, and then incubated with secondary antibody for 1 hour at room temperature. Sections were covered with DAPI-containing mounting media. Samples were imaged using an Evos M5000 (Thermo Scientific, Waltham, MA). IF images were reconstructed using 4X amplification field pictures.

The IF images and their corresponding DGAT-imputed protein spatial plots were spatially registered and adjusted to identical dimensions to standardize scale; matching anatomical regions were manually aligned. All image processing and presence-area assessments were performed using ImageJ (Version 2.0.0-rc-59/1.51n). IF images and their corresponding prediction maps were analyzed to estimate the average ratio between predicted and IF-detected protein-presence areas, stratified by marker and organ. For quantification, the relevant marker channel was isolated (green for ACTA2/αSMA or VIM; red for PECAM1/CD31). For each image pair, three anatomically matched regions of interest (ROIs) with high marker presence were manually selected. The ratio of predicted to observed presence areas was calculated for each ROI. For each sample, the mean of the three ROI ratios represented the predictive accuracy. Sample-level ratios were compiled in GraphPad Prism (v10.2.3). To test for systematic over- or under-prediction, a one-sample t-test was performed for each group against a theoretical ratio of 1.0 (perfect agreement). Statistical significance was set at p < 0.05; p > 0.05 indicated no systematic bias.

### Hardware & Reproducibility

All experiments were conducted on NVIDIA l40s GPUs with 40GB memory. Training a single DGAT model took ∼1 hour for 6 samples. Code and pretrained models are available at: https://github.com/osmanbeyoglulab/DGAT/. Software dependencies are Python 3.10.8, PyTorch 2.0.1, Scanpy 1.11.1, NumPy 1.24.1, SciPy 1.15.3, scikit-learn 1.2.

## Data Availability

The published human tonsil, breast cancer and glioblastoma Visium CytAssist Gene and Protein Expression samples that supports the finding of this study can be downloaded from the 10× Genomics website (https://www.10xgenomics.com/datasets/visium-cytassist-gene-and-protein-expression-library-of-human-tonsil-with-add-on-antibodies-h-e-6-5-mm-ffpe-2-standard; https://www.10xgenomics.com/datasets/gene-and-protein-expression-library-of-human-glioblastoma-cytassist-ffpe-2-standard; https://www.10xgenomics.com/datasets/gene-and-protein-expression-library-of-human-breast-cancer-cytassist-ffpe-2-standard; https://www.10xgenomics.com/datasets/gene-protein-expression-library-of-human-tonsil-cytassist-ffpe-2-standard). The in-house malignant mesothelioma Visium CytAssist Gene and Protein Expression samples and kidney and lung spatial transcriptomics samples used for validation are available in GEO GSE300851. The ST TNBC breast cancer data from Wu et al.^38^, including pathological annotations, were accessed through the Zenodo data repository (https://doi.org/10.5281/zenodo.4739739). The published melanoma ST data that supports the finding of this study can be downloaded from http://www.spatialtranscriptomicsresearch.org/. Additionally, raw count matrices, images, and spatial data for prostate, breast cancer, and lymph node 10x Visium datasets are accessible at https://support.10xgenomics.com/spatial-gene-expression/datasets.

## Acknowledgments

We thank Xiaojun Ma and Annalisa Barrata for helpful feedback on the manuscript. This work was supported by the National Institutes of Health (R35GM146989 and R21CA294196) and by the Centers for Disease Control and Prevention, in association with the National Institute for Occupational Safety and Health and the National Mesothelioma Virtual Bank (U24OH009077). Computational analyses were supported by the University of Pittsburgh Center for Research Computing and the Extreme Science and Engineering Discovery Environment through the Bridges-2 system at the Pittsburgh Supercomputing Center. Histology sectioning was performed by the Pitt Biospecimen Core, and 10x Visium library preparation and Illumina sequencing were conducted by the Health Sciences Sequencing Core at UPMC Children’s Hospital of Pittsburgh. Additional support was provided by the University of Pittsburgh, the Office of the Senior Vice Chancellor for Health Sciences, the Department of Pediatrics, and the Richard King Mellon Foundation for Pediatric Research.

## Author contributions

HW performed computational experiments and analyses, helped to develop the algorithmic approaches and helped to write the paper. BC conducted the pathological review of mesothelioma samples and performed germinal center annotations for tonsil samples. MS performed immunohistochemistry experiments and analyses. LAPF collected and processed the samples and helped with immunohistochemistry analyses. RMF collected and processed the samples, and helped with immunohistochemistry analyses, ASG supervised the experiments and analyses. HUO supervised the project, helped to develop the algorithmic approaches, performed computational experiments and wrote the paper.

